# Adaptive evolution of transcription factor binding affinities

**DOI:** 10.1101/118455

**Authors:** Jungeui Hong, Nathan Brandt, Ally Yang, Tim Hughes, David Gresham

## Abstract

Understanding the molecular basis of gene expression evolution is a central problem in evolutionary biology. However, connecting changes in gene expression to increased fitness, and identifying the functional basis of those changes, remains challenging. To study adaptive evolution of gene expression in real time, we performed long term experimental evolution (LTEE) of *Saccharomyces cerevisiae* (budding yeast) in ammonium-limited chemostats. Following several hundred generations of continuous selection we found significant divergence of nitrogen-responsive gene expression in lineages with increased fitness. In multiple independent lineages we found repeated selection for non-synonymous mutations in the zinc finger DNA binding domain of the activating transcription factor (TF), GAT1, that operates within incoherent feedforward loops to control expression of the nitrogen catabolite repression (NCR) regulon. Missense mutations in the DNA binding domain of GAT1 reduce its binding affinity for the GATAA consensus sequence in a promoter-specific manner, resulting in increased expression of ammonium permease genes via both direct and indirect effects, thereby conferring increased fitness. We find that altered transcriptional output of the NCR regulon results in antagonistic pleiotropy in alternate environments and that the DNA binding domain of GAT1 is subject to purifying selection in natural populations. Our study shows that adaptive evolution of gene expression can entail tuning expression output by quantitative changes in TF binding affinities while maintaining the overall topology of a gene regulatory network.

---

Gene expression evolution is a pervasive source of phenotypic diversity^1–3^. Most genetic variation causing such diversity arises from either *cis*-regulatory or *trans*-regulatory changes. The relative importance of these two mechanisms is the source of a long-standing debate in evolutionary biology^4–7^. Comparative genomics is typically used for inferring the evolutionary processes underlying adaptive evolution of gene expression using extant organisms. However, this approach is limited by the inability to observe the dynamics of gene expression evolution and the challenge of distinguishing neutral from adaptive alleles. LTEE provides a means of performing replicated evolution experiments in controlled conditions and assessing the fitness and phenotypes of evolved lineages and populations. The advent of next-generation DNA sequencing has enabled the comprehensive identification of adaptive alleles and their dynamics during LTEE^8–11^. In LTEE performed in conditions of constant nutrient-limitation using chemostats, the expression of nutrient transporter genes is a primary target of selection^12^. Copy number variants (CNVs), comprising gene amplifications that result in increased expression, are a recurrent class of adaptive alleles in chemostat selections. In addition, adaptive mutations in *trans* factors such as transcription factors (TFs) are frequently identified in LTEEs^8,11^. These variants are primarily missense, frame-shifting or stop codon mutations that are likely to confer strong effects on gene expression in evolved lineages^13,14^. However, how protein coding changes in transcriptional regulators identified during LTEE alter gene expression and confer increased fitness remains unknown.

Previously, we identified repeated selection within a single LTEE population maintained in ammonium-limited chemostats of independent missense mutations in the DNA binding domain of GAT1^11^. *GAT1* encodes a transcriptional activator of nitrogen catabolite repression (NCR) genes, which encode proteins required for uptake and utilization of multiple nitrogen sources in *Saccharomyces cerevisiae.* The expression of NCR genes is regulated by four GATA factors – two activators (GAT1 and GLN3) and two repressors (DAL80 and GZF3) (**Supplementary Fig. 1**) – that bind the same GATAA binding sites in promoter regions^15^. NCR is an ideal system for studying the evolution of gene expression owing to the well-characterized properties of its four regulators and small number of direct targets (~ 41). In this study, we aimed to determine whether adaptive mutations in *GAT1* result in alteration of its regulatory activities and, if so, how these changes impact fitness in different conditions.

First, we tested the repeatability of adaptive *GAT1* mutations in initially clonal populations maintained in ammonium-limited environments by performing LTEE in triplicate using asexually reproducing populations in ammonium-limited chemostats (Fig. 1a). We found that evolution is highly parallel both at the phenotypic and genotypic levels over 250 generations. We observed significant increases in population level fitness with an overall deceleration in the rate of fitness improvement with time as seen in previous LTEEs^16^ (Fig. 1b). Using whole genome, whole population sequencing of evolving populations, we identified recurrent selection for missense mutations in the DNA binding domain of *GAT1* during the early stages of adaptive evolution. Consistent with positive selection for increased ammonium transport capacity, we also identified CNVs that include *MEP2*, which encodes a high-affinity ammonium transporter, at high frequencies following 250 generations of selection (Fig. 1c).

**Figure 1.**
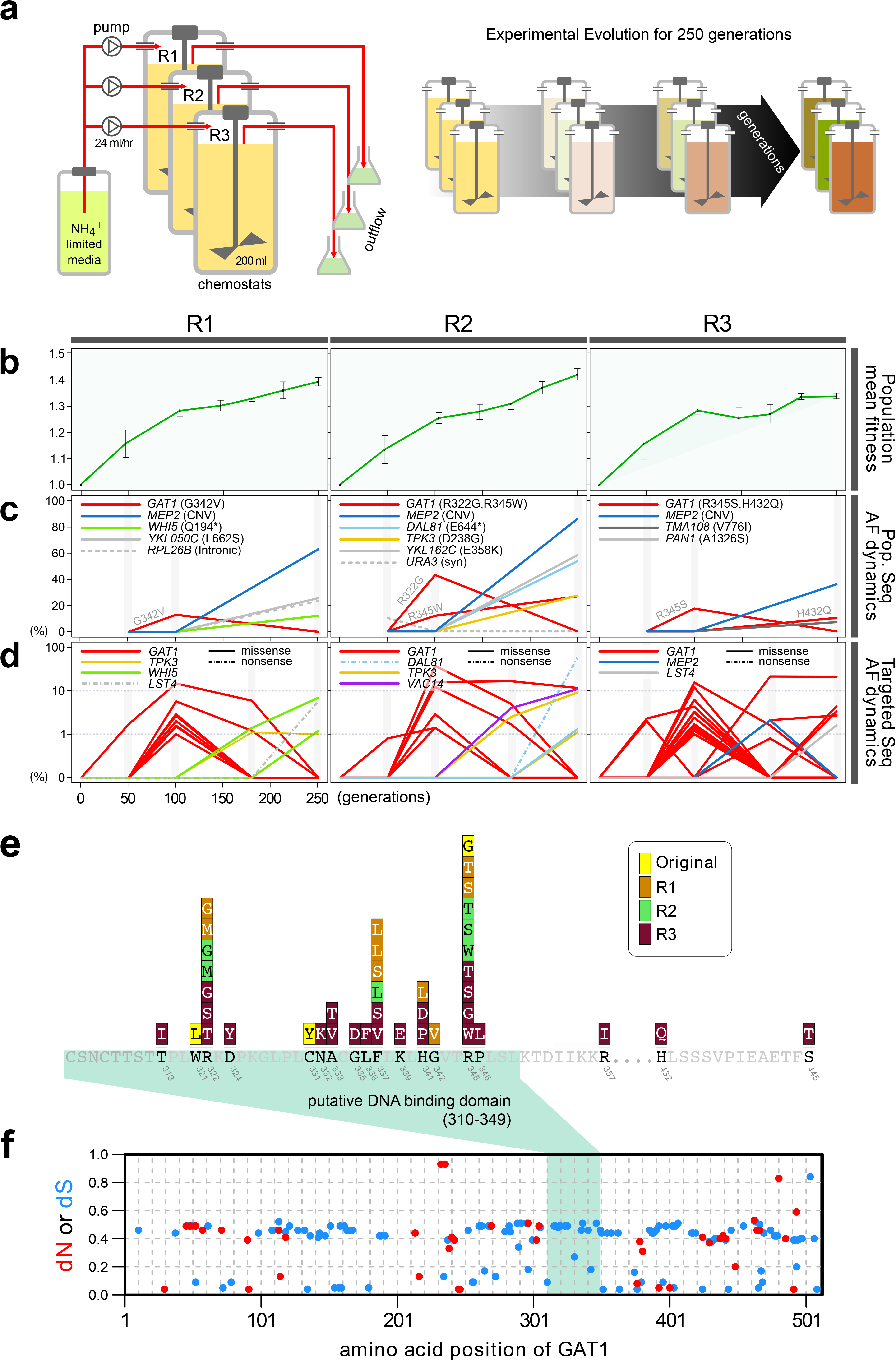
Repeated LTEE in ammonium-limited chemostats selects for adaptive mutations in *GAT1.* **a, LTEE design.** Three chemostat vessels were founded with a clonal ancestral population and maintained under constant dilution with defined media containing 0.4mM (NH_4_)_2_SO_4_, (0.8mM nitrogen). Chemostats were maintained in continuous mode for 250 generations (~ 3 months) and samples obtained every 10-20 generations for archiving. **b, Dynamics of population fitness.** The rate of fitness improvement decelerates over time for three independent replicated LTEE (R1, R2 and R3). Error bar are 95% CI of linear regression analysis of competition assays, which comprised six time points each. **c, Allele dynamics in parallel LTEE**. Whole population while genome sequencing was performed on samples from the three LTEEs at 50, 100 and 250 generations to identify high frequency mutations (> ~10%) using an Illumina HiSeq2500 in 2×50bp paired end mode with an average read depth of ~ 50X. The frequency of the *MEP2* amplification was defined as the proportion of clones bearing more than 2 copies of *MEP2* among 96 randomly selected clones. *GAT1* (red lines) variants are a primary target in the earliest generations of selection but, in some LTEE, are ultimately replaced by other alleles including CNVs that encompass *MEP2* (blue lines) and mutations in genes that control cell cycle and growth. **d, Dynamics of minor frequency mutations.** 12 genes that have previously been identified as adaptive targets of selection in different nitrogen-limited environments (see method) were subjected to targeted deep sequencing. Multiple missense mutations in *GAT1* are present simultaneously in each population and compete with each other during the earliest generations. **e. Mutational landscape of adaptive** *GAT1* **mutations.** All mutations in *GAT1* with population AF greater than 1% are shown. The DNA binding domain of GAT1 is a mutational hotspot in all ammonium-limited LTEE. Multiple missense (protein coding alteration) mutations were observed but no nonsense or frame-shift mutations were detected. **f, The DNA binding domain of GAT1 is under purifying selection in the wild.** *dN* and *dS* values for GAT1 at each amino acid position was calculated using SNAP v1.1.1 (http://www.hiv.lanl.gov/content/sequence/SNAP/SNAP.html) using sequences from 42 different wild yeast strains. Unlike LTEE in ammonium-limited chemostats, the *GAT1* DNA binding domain is under purifying selection implying that non-synonymous mutations are likely to be detrimental in dynamic environments.

Recently, lineage tracking during LTEE in large populations has shown that many independently derived beneficial alleles are present at low frequencies and unlikely to be detected using whole genome, whole population sequencing^17^. Therefore, we investigated whether additional low frequency mutations of *GAT1* were present in evolving populations using targeted ultra-deep sequencing (Fig. 1d and **Table S1**; see **methods**). Evolving populations contain multiple different *GAT1* mutations at frequencies of 10^−2^–10^−3^ that are most abundant during the earliest stages of adaptive evolution before lineages containing *MEP2* CNVs begin to dominate the populations. Importantly, all mutations are missense mutations in the DNA binding domain of GAT1 that we find is under strong purifying selection in natural environments (Fig. 1e and 1f).

To study the functional basis of adaptive *GAT1* mutations, we isolated three individual alleles from distinct evolved lineages – *gat1-1* (W321L), *gat1-2* (C331Y), and *gat1-3* (R345G) – (Fig 2a) using backcrossing and allele specific PCR genotyping^11^. Using competition assays (see **methods**), we determined the fitness of the evolved lineages that contain *GAT1* mutations plus an additional 3-4 variants in other genes, and of the *GAT1* mutations alone, in different nitrogen-limiting conditions – ammonium, glutamine, proline and urea – using chemostats in addition to rich media (YPD) batch condition. All strains containing *GAT1* missense mutations are significantly increased in fitness in ammonium-limited chemostats, but exhibit dramatically decreased fitness in all other nitrogen-limited conditions consistent with antagonistic pleiotropy (Fig. 2b). By contrast, a *GAT1* knock out (KO) strain exhibits a fitness defect in both ammonium and proline limited chemostats. *GAT1* mutations are nearly neutral in YPD batch media conditions in which *GAT1*, and the entire NCR regulon, is transcriptionally repressed. Collectively, these results suggest that missense mutations in the DNA binding domain of GAT1 alter its function rather than rendering the TF non-functional.

**Figure 2.**
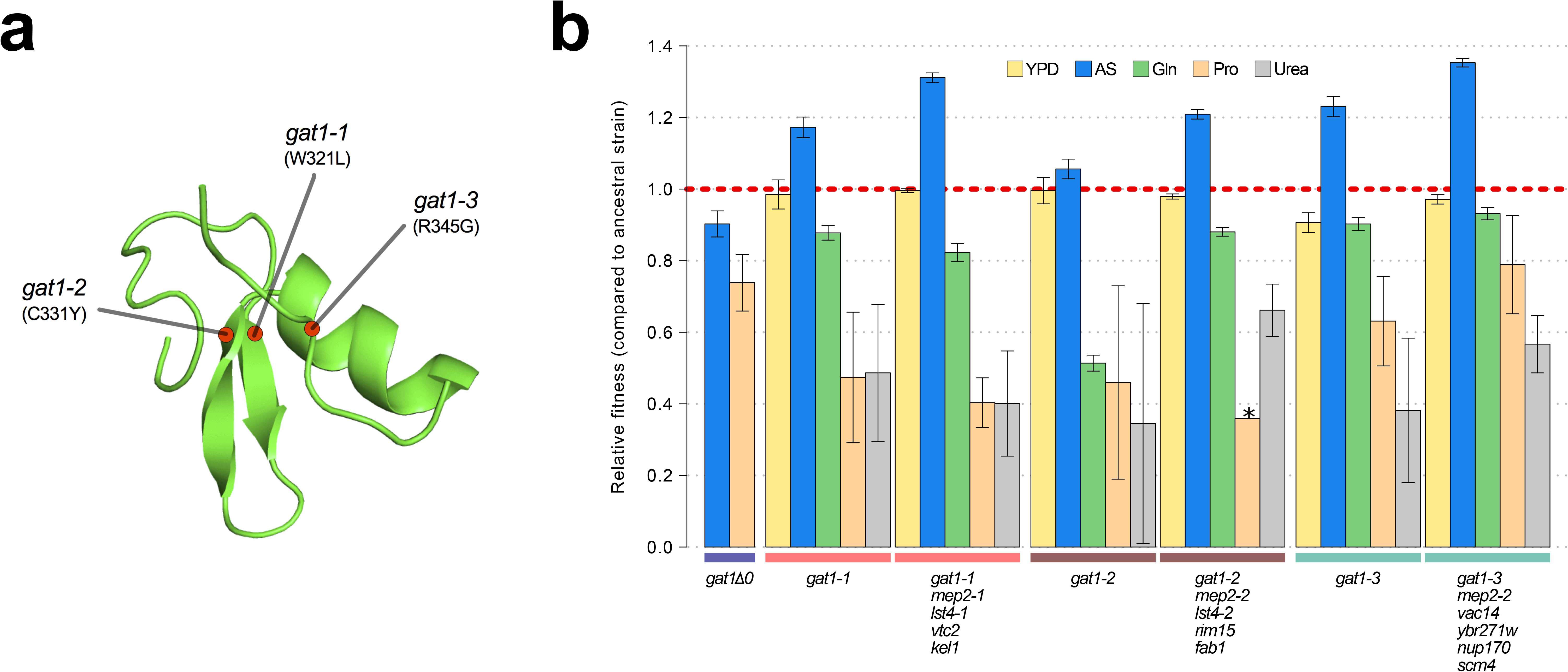
Adaptive *GAT1* mutations confer antagonistic pleiotropy. **a, 3D structure model of GAT1 DNA binding domain.** The 3D structure is based on available GATA factor DNA binding domain structures in the modebase database (https://modbase.compbio.ucsf.edu/). Adaptive amino acid changes occur in the functional domain of GAT1. **b**. **Fitness effects of *GAT1* mutations. *GAT1*** variants are beneficial in ammonium-limited chemostats and confer increased fitness in the lineages in which they occur. *GAT1* variants confer a fitness cost in non-ammonium limited environments consistent with antagonistic pleiotropy. Error bars represent 95% CI of linear regression analysis of competitive fitness assays. (YPD : YPD batch culture, AS : Ammonium sulfate-limited chemostat, Gln : Glutamine-limited chemostat, Pro : Proline-limited chemostat, Urea : Urea-limited chemostat; all nitrogen-limited media were normalized to 0.8mM nitrogen).

Increased gene expression of nutrient transporters through gene amplification is a dominant mechanism for increased fitness in nutrient-limited chemostats^11^. We asked whether the effects of beneficial *GAT1* mutations are mediated by their impact on *MEP2* regulation and therefore represent an alternate class of adaptive mutations with the same functional effect as CNVs. To test the importance of *MEP2* for the increased fitness associated with *GAT1* variants we deleted either the ORF or the promoter region (1kb upstream of the ORF) of *MEP2* in the background of the *gat1-1* and *gat1-3* alleles and tested their fitness in ammonium-limited chemostats. All strains have severely reduced fitness (**Supplementary Fig. 2**) consistent with the beneficial effect of *GAT1* variants being dependent on *MEP2* expression. Given that both the promoter region and ORF of *MEP2* is required for beneficial effect of *GAT1* adaptive alleles, we inferred that the effect of selected *GAT1* variants is likely to be mediated through altered transcriptional regulation of *MEP2.*

We investigated the effect of adaptive missense mutations in *GAT1* on gene expression. We performed RNAseq using strains that contain each of the three adaptive *GAT1* mutations alone cultured in ammonium-limited chemostats. We compared expression profiles of *GAT1* adaptive mutations with those in the corresponding evolved lineages from which the mutations were isolated^11^. We find significant divergence of NCR gene expression in all tested strains (Fig. 3a). *GAT1* mutations result in increased expression of genes encoding ammonium transporters *(MEP1, MEP2,* and *MEP3)* as well as transporters that import other nitrogen sources such as urea, allantoin and GABA. The evolved lineages, which contain *GAT1* mutations as well as additional mutations, show more “fine-tuned” gene expression in which only ammonium permease-encoding genes are up regulated while other NCR targets, which are likely to be irrelevant when ammonium is the only nitrogen source, are repressed. Interestingly, *GAT1* mutations also result in increased expression of *GAT1* mRNA and reduced expression of *DAL80* mRNA.

**Figure 3.**
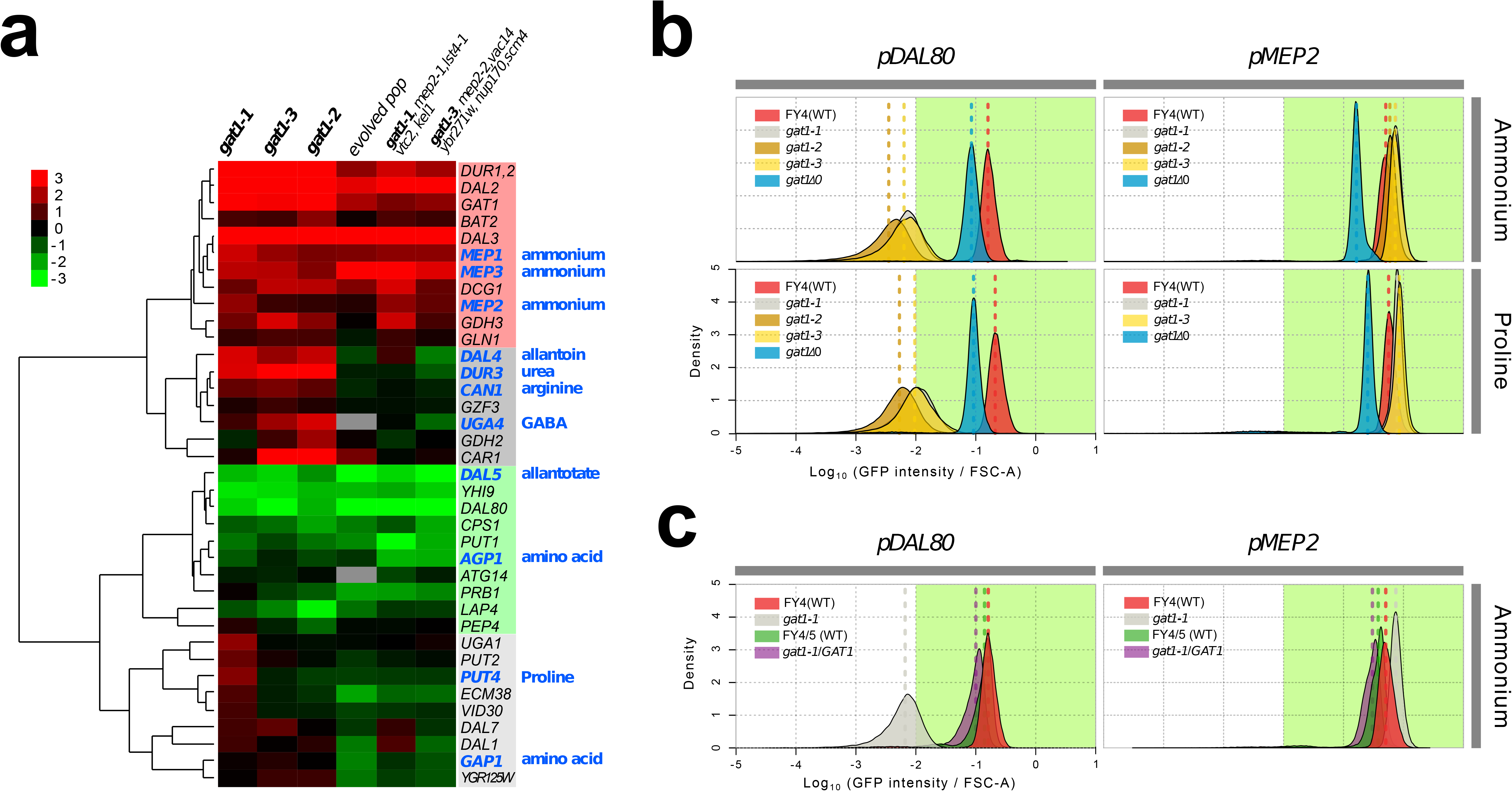
Adaptive *GAT1* variants exhibit differential effects on target gene expression. **a, Gene expression profile of NCR genes in different** *GAT1* **variant lineages.** ‘Blue’ gene names indicate permease-encoding genes for different nitrogen sources. Clustering expression values relative to the ancestral strain are shown for 41 experimentally confirmed NCR target genes. Expression data for the final evolved population and two clonal lineages are from^11^. **b, Transcriptional reporter assay for *MEP2* and *DAL80.*** We quantified GFP expression from the *MEP2* and *DAL80* promoters in the ancestral and *GAT1* variant backgrounds in different nitrogen-limited conditions. In nitrogen-limited chemostats containing different individual sources of nitrogen (ammonium, glutamine, proline and urea), all mutant strains *(gat1-1, gat1-2* and *gat1-3)* show the same gene expression pattern for *DAL80* (suppressed) and *MEP2* (activated) compared to the ancestral genotype. A *GAT1* KO mutant expresses both *DAL80* and *MEP2* to a significantly reduced degree. **c. GFP reporter assay for *MEP2* and *DAL80* promoters in heterozygous diploids**. The *gat1-1/GAT1* genotype results in expression of *DAL80* or *MEP2* comparable to the ancestral strain.

To quantify changes in transcriptional activity attributable to *GAT1* variants, we fused the promoter sequences of four targets of GAT1 (*GAP1, MEP2, GZF3* and *DAL80*) to the coding sequence of GFP in the backgrounds of the ancestral (WT), KO (*gat1 A0*), and the three GAT1 variants (*gat1-1, gat1-2* and *gat1-3*) (see **Supplementary Fig. 3a**). GFP expression levels measured using this assay were highly comparable to RNAseq data and therefore a good proxy of transcriptional activation by the different GAT1 variants of each promoter (**Supplementary Fig. 3b** and **3c**). Using these reporter constructs, we studied transcriptional activation by the different variants of GAT1 in a variety of conditions.

We found significant differences in transcriptional activities between *GAT1* variants at *MEP2* and *DAL80* promoters in nitrogen-limited (both ammonium and proline) conditions (Fig. 3b). All adaptive *GAT1* mutations resulted in reduced *DAL80* expression and increased *MEP2* expression compared to the wild type, while a *GAT1* KO strain showed the opposite pattern. This result is consistent with our RNAseq analysis and provides additional evidence that the beneficial missense mutations in the DNA binding domain of GAT1 do not render it nonfunctional. We also tested whether these mutations are dominant or recessive using diploid strains heterozygous for adaptive *GAT1* mutations (e.g. *gat1-1/GAT1)* (Fig. 3c). *DAL80* and *MEP2* promoter activities in these heterozygotes was identical to the haploid ancestral *GAT1* strain indicating that the missense mutations of *GAT1* are recessive and therefore unlikely to have gain-of-function or dominant negative effects.

To test whether adaptive *GAT1* mutations have acquired new binding specificities we used protein binding microarrays (PBMs) to assay the DNA binding domains of adaptive *GAT1* alleles *(gat1-1* and *gat1-3)* as well as the ancestral *GAT1.* Whereas ancestral *GAT1* shows clear evidence of specificity for the GATAA consensus sequence, adaptive *GAT1* alleles failed to show evidence for any significant binding specificity using PBMs (**Table S2**) indicating that their functional and fitness effects are not exerted by acquisition of new DNA binding specificities. We quantified alterations in the affinity of the adaptive GAT1 alleles for their target DNA sequences using electrophoretic mobility shift assays (EMSAs) (Fig. 4a, **Table S3** and **S4**). We studied the binding of adaptive GAT1 alleles to the promoter sequences of *MEP2* and *DAL80*, which showed discrepant effects of adaptive GAT1 alleles based on transcriptional reporters (Fig. 3b) and RNAseq (Fig. 3a). *MEP2* contains two distinct GATAA sequence motifs in its promoter region whereas *DAL80* possesses only one. Adaptive *GAT1* variants showed significantly decreased binding affinity to all target motifs (Fig. 4b). Binding kinetics calculated based on a two parameter Michaelis-Menten model^18^ show that adaptive *GAT1* mutations have a more detrimental effect on binding to the *DAL80* promoter GATAA site compared with the two *MEP2* promoter binding sites likely reflecting differences in motif strength (Fig. 4c and **Table S5**). This suggests that the adaptive *GAT1* alleles still encode functional TFs that possess decreased binding affinity to their targets compared to the ancestral allele. Whereas reduced affinity of GAT1 variants for the comparatively weak promoter of *DAL80* results in its reduced expression, activation of the comparatively strong *MEP2* promoter by GAT1 variants may be minimally perturbed by a reduction in GAT1 affinity. Consistent with a differential effect of reduced GAT1 affinity dependent on the strength of individual promoters, we identified a significant positive correlation between gene expression levels of NCR genes and the estimated affinity of each promoter for GAT1 (Fig. 4d). Thus, while the overall topology of NCR regulation is maintained, the increase in *MEP2* expression, and potentially other targets with strong promoters (including *GAT1* itself), is mediated by both maintenance of direct binding, resulting in direct activation, and an indirect effect through reduced activation of the repressor *DAL80* (Fig. 4e).

**Figure 4.**
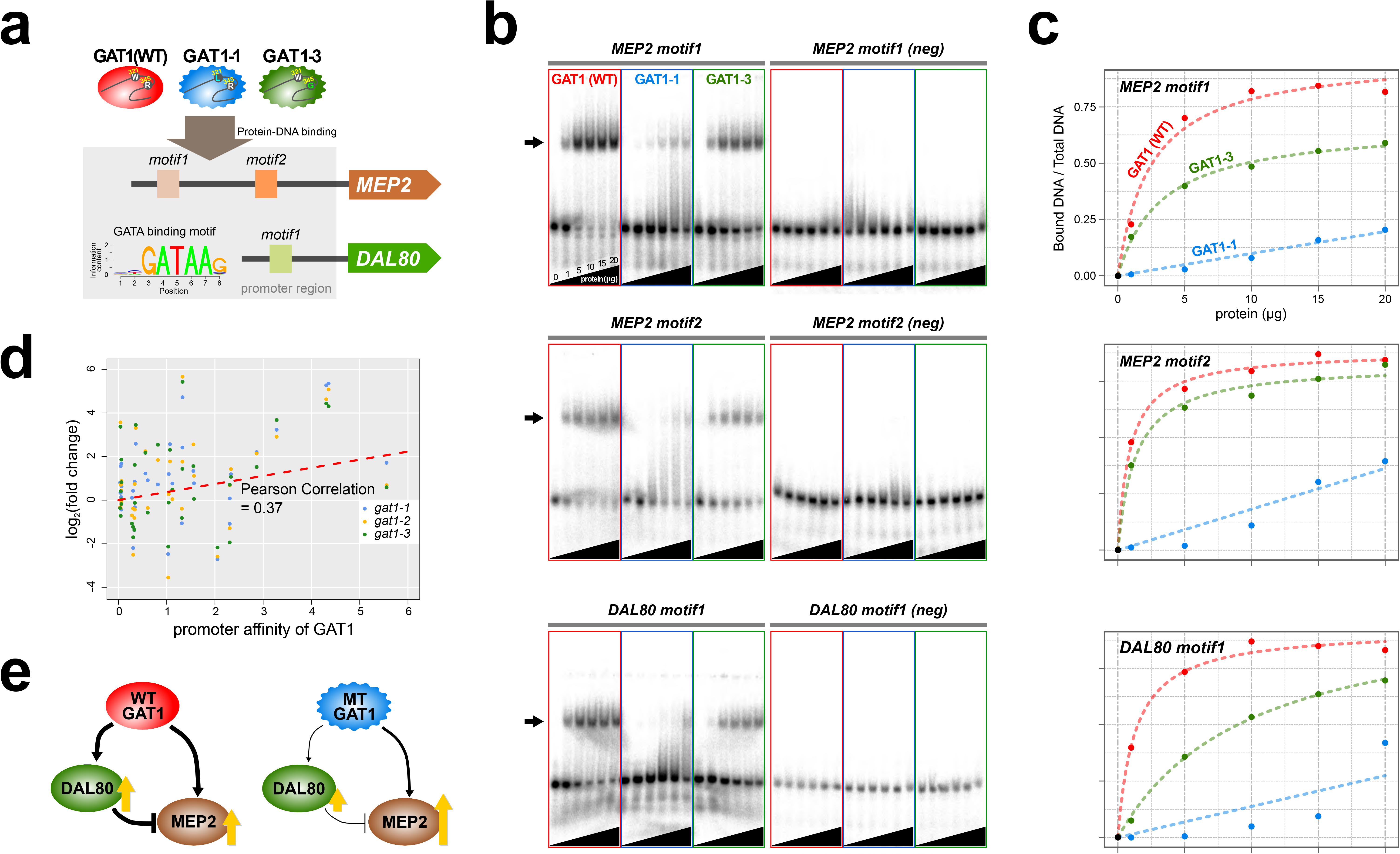
Adaptive *GAT1* variants alter DNA binding affinities in a promoter-specific manner. **a, Design of protein-DNA binding assays.** We purified a 151 amino acid long protein fragments of *GAT1* containing the DNA binding domain (DBD) (positions 310-349) for the ancestral *GAT1* and two adaptive mutations – *GAT1-1* (W321L) and *GAT1-3* (R345G). DNA sequences containing the consensus sequence in *MEP2* and *DAL80* promoters were determined from the MotifDb collection in Bioconductor. **B, Electrophoretic mobility shift assay (EMSA) analysis of *GAT1* variants.** Increasing quantities of the GAT1 DBD protein fragment from 0-20μg were added to ^32^P-labeled dsDNA. Arrows and stars indicate protein-bound DNA and unbound DNA respectively. Negative motif sequences were prepared by randomizing bases only at their GATAA binding motif sequences (see **Table S4**). **c**, **Proportion of bound DNA for different amounts of protein**. The two GAT1 variants showed significantly decreased binding affinity to all three GATAA motifs. Dotted lines are fits to a two-parameter Michaelis-Menten (MM) model. Only ancestral *GAT1* and *GAT1-3* (R345G) showed significant estimates of two parameters *(Fr_max_* and *K_x_)* (p-value < 0.02). Insignificant estimates for *GAT1-1* (W321L) may be due to the small number of data points (**Table S5**). **d, Correlation between GAT1 promoter affinity and expression changes for NCR target genes**. The promoter affinity of GAT1^24^ is positively correlated (Pearson correlation = 0.37; p-value = 0.02) with the alteration in expression of NCR targets attributable in strains containing adaptive *GAT1* mutations (log_2_ transformed fold change of gene expression of NCR genes in *GAT1* variant strains compared to the ancestor strain). **e, Model for altered output from the NCR regulatory network by adaptive *GAT1* mutations**. *GAT1* mutations reduce affinity for the promoter of *MEP2*, but have an even greater reduction in affinity for the promoter of the repressor, *DAL80.* The simultaneous maintenance of MEP2 activation by GAT1 variants and decreased suppression of *MEP2* through reduced abundance of DAL80 results in a net increase in *MEP2* expression in evolved lineages compared to the ancestral strain.

Here, we describe the evolution of a transcriptional circuit in real time and dissection of its functional effects. Missense mutations in *GAT1* are under strong positive selection in conditions of ammonium-limitation. The functional effects of *GAT1* mutations are exerted by tuning the output of an incoherent feed-forward loop through alteration in the binding affinities, but not specificity, of the TF. The repeated evolutionary dynamics suggest that remodeling the transcriptional regulatory network is a dominant and repeatable mode of adaptive evolution during the earliest stages of evolution but might not represent a fitness optimum as seen in the mutually exclusive dynamics pattern between missense mutations in GAT1 and amplification of *MEP2* in the Fig. 1c.

Experimental evolution in eukaryotic microbes has many parallels with the molecular processes, and evolutionary dynamics, underlying tumorigenesis. The evolutionary hotspot in the DNA binding domain of *GAT1* identified in this study is reminiscent of hotspots in transcriptional regulators important in tumorigenesis including recurrent missense mutations in the DNA binding domain of TP53, a driver in the majority of human cancers^19^, and CTCF, a poly-zinc finger transcription factor regulating oncogenes and tumor suppressor genes^20^. As with adaptive evolution, missense mutations in the DNA binding domain of these transcription factors result in altered binding affinity to the promoters of target genes resulting in aberrant transcriptional activation in various cancers^21–23^. Our study highlights the importance of incorporating biophysical parameters in gene regulatory networks models in order to predict their functional output in the context of both evolution and disease.

## METHODs SUMMARY

LTEE was performed using clonal populations founded with an ancestral haploid strain (FY4, isogenic to S288c) cultured in 0.8mM ammonium-limited media in chemostats at a constant dilution rate (0.12 hr^−1^) for 250 generations (2 months) during which whole population samples were regularly archived. We identified the comprehensive set of adaptive alleles including SNPs, indels and CNVs and determined their allele frequencies and dynamics in evolving populations using Illumina sequencing and qPCR. Competition assays were conducted with the ancestral strain to infer the relative fitness of evolved lineages in identical nitrogen-limited chemostats. Purifying selection on the DNA binding domain of GAT 1 in natural yeast isolates was confirmed by a dN/dS test. Directional RNA-seq and GFP reporter assay were used to identify alterations in the expression of the NCR regulon due to adaptive GAT1 alleles. Using *in vitro* protein binding microarrays (PBM) and electrophoretic mobility shift assays (EMSA), we quantified alterations in the affinity of the GAT1 DNA binding domain to the promoter of two of its primary targets, *MEP2* and *DAL80*.

